# Pre-stimulus beta-band activity is a signature of statistical learning

**DOI:** 10.1101/2020.04.15.042507

**Authors:** Louisa Bogaerts, Craig G. Richter, Ayelet N. Landau, Ram Frost

## Abstract

Statistical learning (SL) is taken to be the main mechanism by which cognitive systems discover the underlying regularities of the environment. We document, in the context of a classical visual SL task, divergent rhythmic EEG activity during the anticipation of stimuli within patterns versus pattern transitions. Our findings reveal differential pre-stimulus oscillatory activity in the beta band (∼20 Hz) that indexes learning: it emerges with increased pattern repetitions, and importantly, it is highly correlated with behavioral learning outcomes. These findings hold the promise of converging on an online measure of learning regularities and provide important theoretical insights regarding the mechanisms of SL and prediction.

**Significance Statement:** SL has become a major theoretical construct in cognitive science, providing the primary means by which organisms learn about regularities in the environment. As such it is a critical building block for basic and higher-order cognitive functions.

Here we identify for the first time a spectral neural index in the time window prior to stimulus presentation, which evolves with increased pattern exposure, and is predictive of learning performance.

The manifestation of learning that is revealed not in stimulus processing but in anticipatory moments of the learning episode, makes a direct link between the fields of statistical learning and predictive processing, and suggests a possible mechanistic account of visual SL.

## Introduction

Sensory information is oftentimes structured, both in time and in space. *Statistical learning* (SL) is a cognitive mechanism by which we incidentally discover regularities in the environment. Seminal work by Saffran and colleagues demonstrated that infants can extract syllable patterns presented in a continuous speech stream based solely on the extent of the syllables’ co-occurrences (Saffran et al., 1996). Ever since, SL has been extensively investigated and was documented in individuals of all ages, over different types of stimuli and sensory modalities (see Frost, Armstrong, & Christiansen, 2019, for a recent review). Similar to linguistic information, our visual environment contains extensive statistical regularities (e.g., probabilistic relations between objects, prevalent sequences of letters, etc.). Indeed, recent studies of *visual* SL demonstrated robust learning of temporal relationships among sequentially presented ordered stimuli (Bogaerts, Siegelman, & Frost, 2016; Fiser & Aslin, 2002; Kirkham, Slemmer, & Johnson, 2002). SL is unsupervised and may occur without intent or awareness of the structured nature of the input (Aslin et al., 1998; Fiser & Aslin, 2001; Turk-Browne et al., 2005).

While SL has become an important construct in cognitive science, little is known about the neural mechanisms underlying it. Most neuroimaging studies that identified brain regions sensitive to the statistical regularities in sensory input, have summarized brain activity during structured vs. unstructured blocks, lasting several seconds or minutes. These studies associated SL effects with domain-general regions involved in binding temporal and spatial contingencies such as the hippocampus, the medial temporal lobe and inferior frontal gyrus (e.g., Karuza et al., 2013; Shohamy & Turk-Browne, 2013; Turk-Browne, Scholl, Chun, & Johnson, 2009). Investigating item-specific hemodynamic responses, Turk-Browne and colleagues (2010) also reported increased hippocampal activity in response to visual stimuli which predict subsequent visual stimuli. At the same time, imaging work identified regions in the early visual and auditory cortices that are sensitive to regularities in vision and audition (e.g., McNealy, Mazziotta, & Dapretto, 2006; Turk-Browne et al., 2009; see Frost, Armstrong, Siegelman, & Christiansen, 2015, for discussion).

Here we aimed to go beyond the neurobiological “where in the brain” of SL, focusing instead on underlying neurobiological mechanisms. Targeting visual SL, we investigated whether brain rhythms play a role in the implicit learning of statistical regularities. So far, attempts to identify neural indices of SL with electrophysiology centered on the consequences that learning has on stimulus processing. The consequences of learning were typically assessed by contrasting evoked responses to stimuli which are predictable to those of stimuli which can not be predicted (*N400* in the visual domain: Abla & Okanoya, 2009; *N400*, *N100*, *P300* and *P200* in the auditory domain: e.g., Abla, Katahira, & Okanoya, 2008; Batterink, Reber, Neville, & Paller, 2015; Cunillera, Toro, Sebastián-Gallés, & Rodríguez-Fornells, 2006; De Diego Balaguer, Toro, Rodriguez-Fornells, & Bachoud-Lévi, 2007; Sanders, Newport, & Neville, 2002; and see Daikoku, 2018 for a recent review). Abla and Okanoya (2009), for example, demonstrated that participants who show behavioral evidence for good learning have an increased *N400* in response to unpredictable shapes in a sequence, compared to the middle and final shapes within a sequence, which are fully predictable. ERPs were also shown to differentiate between visual stimuli that carry high vs. low predictability for a subsequent target. Highly reliable predictors were shown to elicit either a larger centro-parietal late positivity (Daltrozzo et al., 2017) or a larger P300 component (Jost et al., 2015). More recently the predictability of visual stimuli was shown to modulate the low-frequency activity associated with stimulus presentation: in comparison to predictable second items of a learned pair, unexpected shapes elicited stronger activity in the alpha (7 – 14 Hz) range (Zhou et al., 2019). Auditory SL has also been indexed considering longer segments of structured vs. unstructured input, by quantifying the neural entrainment to the temporal structure created by repeating regular sequences (Batterink & Paller, 2017; Farthouat et al., 2017). This entrainment is postulated to reflect the perceptual binding of stimuli into familiar composites (Batterink & Paller, 2017).

In the present study by contrast, our main interest was in EEG activity in the time window leading up to stimulus presentation, targeting the mechanisms of prediction. Underlying our approach are two notions on the nature of SL. First, the learning process itself is likely not a passive process and therefore the online manifestations of learning need not be limited to stimulus processing but could impact anticipatory moments in the learning episode. Second, the outcome of learning, once again, might entail an active state of anticipation whereby learned regularities lead to active predictions (e.g., Engel, Fries, & Singer, 2001; Turk-Browne et al., 2010; see also Tollman, 1932). In previous studies on SL, the process of prediction has often been implied but not operationally measured (e.g., Abla & Okanoya, 2009; Sanders et al., 2002). Our present investigation uses EEG to focus on the pre-stimulus epoch in the absence of stimulus processing, aiming to measure anticipation directly. Importantly, as we are not focusing on stimulus evoked responses, analyzing pre-stimulus epochs enables us to quantify spectral signatures of brain activity – i.e., brain rhythms.

Indeed, experimental work with animals as well as humans has demonstrated that rhythmic brain activity from the delta to gamma range (1-100 Hz) has functional relevance for several sensory and cognitive processes (Buzsáki, 2016). For example, activity and oscillatory synchrony in the Beta frequency (13 – 30 Hz) has been associated with top-down modulation (e.g., Hipp, Engel, & Siegel, 2011; see Bressler & Richter, 2015, for a review). In addition, theta-band activity (4 – 7 Hz) has been associated with the categorical prediction of upcoming images (e.g., Cashdollar, Ruhnau, Weisz, & Hasson, 2017), and locations in visual-search displays (e.g., Spaak & de Lange, under review).

Here, we document, for the first time, divergent rhythmic pre-stimulus EEG activity within patterns versus between patterns (i.e. pattern transitions). We demonstrate –in the context of a typical visual SL task– increased power in the beta band at pattern transitions. We further find that this differential pre-stimulus beta band activity is a signature of learning: we show that it emerges with increased pattern repetitions, and importantly, we show that it is highly correlated with behavioral learning outcomes.

## Materials and Methods

### Participants

Thirty-five participants of the Hebrew University (21 females) participated in the study for payment or for course credit. Participants had a mean age of 24.8 years (range = 17-33) and reported normal or corrected-to-normal vision and no history of neurological or psychiatric disease. Written informed consent was obtained from all participants in line with the institutional IRB approval from the Hebrew University of Jerusalem.

### Experimental design

The experiment consisted of a structured familiarization stream with embedded patterns, directly followed by a test, and of a random stream. The random stream either followed the test or preceded the structured stream (counterbalanced across participants).

The task included 24 abstract shapes (see Figure 1) in dark orange (R=205, G=85, B=50), displayed on a grey background (R=110, G=110, B=110). The latent structure of the structured visual input stream was similar to that of multiple previously employed SL tasks (e.g. Frost, Siegelman, Narkiss, & Afek, 2013; Glicksohn & Cohen, 2013; Turk-Browne, Junge, & Scholl, 2005). Shapes were randomly organized for each participant to create eight triplets, with a transitional probability (TP) of 1 between shapes within each triplet. The structured stream consisted of 54 repetition blocks, with all eight triplets appearing once (in a random order) in each repetition block, with the constraint that a same triplet could not appear twice in a row. A self-paced break was included after every 6 repetition blocks, dividing the structured familiarization stream into 9 equal periods. Given our interest in anticipatory brain activation we presented stimuli for a short duration (0.2 s) with a fixed interstimulus interval (ISI) of 1.1 s. Prior to exposure to the structured stream participants received the instruction “In this part there are shapes that follow each other, pay attention to the sequence. Following this part you will be asked questions about what you have seen”.

**Figure 1.**
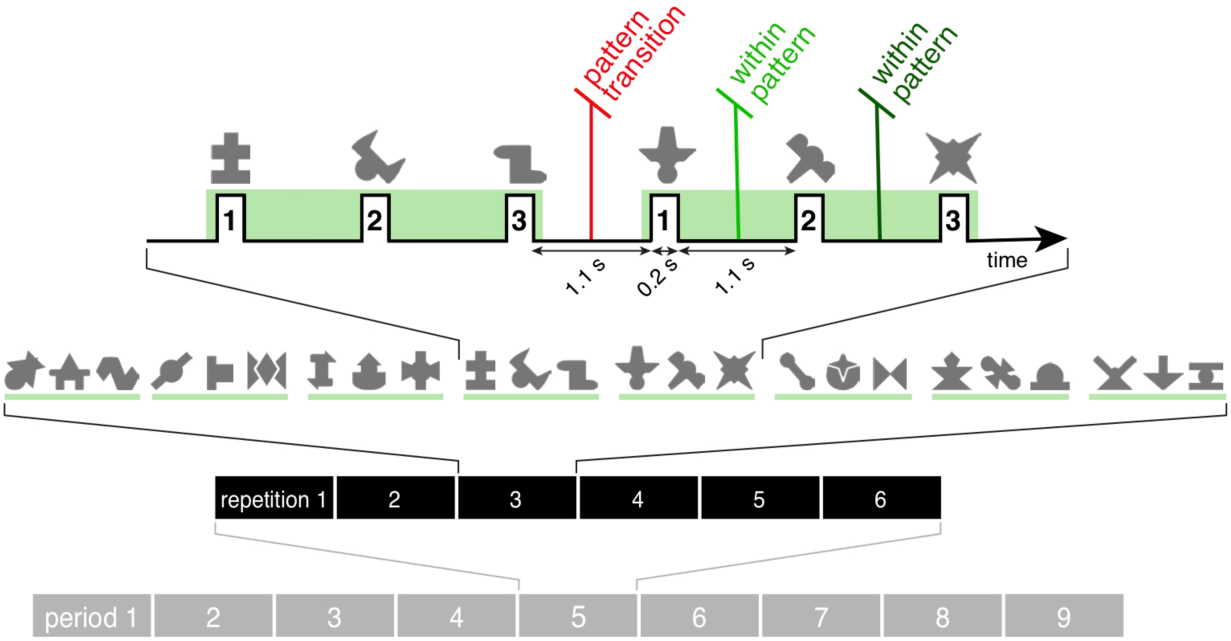
Schematic depiction of the structured familiarization stream, containing 9 exposure periods each consisting of 6 repetitions of 8 embedded triplets.

Following exposure to the structured stream, participants completed an offline test, consisting of 32 two alternative forced choice questions. For each question, participants were instructed to select which of two possible triplet sequences they believed had occurred during the structured familiarization stream. For each question, participants were sequentially presented with: (1) a target – three shapes that formed a triplet in the structured stream, and (2) a “foil” – three shapes that never appeared in sequence in the structured stream. Foils were constructed without violating the position of the shapes within the triplets (e.g., given the triplets ABC, DEF and GHI, a possible foil could be, for example, AEI or GBF but not BID). Shapes making up a target or foil appeared sequentially with a fixed presentation rate and ISI as during exposure. A blank screen of 1.5 s separated the two three-item sequences. The offline test score, defined as the number of correct identifications of targets, ranged from 0 to 32. Chance performance corresponds to a score of 16/32.

The random stream consisted of 18 repetition blocks, with the same 24 shapes appearing once in each repetition block, in a random order (with timing parameters identical to those in the structured stream). Prior to their exposure to the random stream, participants received the instructions “In this part we want to test whether your brain recognizes individual shapes. We will therefore present the same shapes again and again. Your task is simply to watch the shapes attentively.”

### EEG recording

EEG was recorded from 64 Ag/AgCl active electrodes (g.tech system, 62 scalp electrodes and 2 electrode earclips). Four EOG electrodes were placed at the outer canthi of both eyes (horizontal EOG), and above and below the left eye (vertical EOG). Eye position and pupil area were monitored (binocularly) with a video-based eye tracker (Eyelink 1000, SR Research Ltd., Mississauga, Ontario, Canada). A chinrest was used to reduce head movements. We used a Simulink model in Matlab (2015a, The Mathworks Inc., Natick MA) with a sampling rate of 512 Hz as data acquisition software for both the EEG and eye data.

### EEG data preprocessing

All EEG analyses were carried out using the Fieldtrip toolbox (Oostenveld et al., 2011) for Matlab (2016b, The Mathworks Inc., Natick MA). EEG preprocessing was conducted using the following analysis pipeline: 1) Scalp signals were referenced to the average of the left and right earlobes. 2) Signals were bandpass filtered between 0.1 and 140 Hz with 50 and 100 Hz line noise removal. 3) The data was segmented into 1.3 s epochs (0.8 s prior to 0.5 s after shape onset). 4) We identified bad channels (mean number per participant = 1.86, SD = 2.21) and eliminated epochs containing large artifacts, based on an absolute amplitude threshold and a variance threshold using Fieldtrip’s artifact detection routines (mean threshold absolute amplitude = 237.06 µV, SD = 103.30 µV; mean threshold variance = 3381.243 µV, SD = 3231.28 µV). On average, 4.30% of epochs were removed for each participant (range = 0.87% - 12.15%). 5) Large muscle artifacts were identified using automatic artifact detection, and were subsequently visually inspected. Epochs containing confirmed muscle artifacts were rejected (average of 1.78% of epochs rejected, range = 0.23% – 5.79%). 6) Blink artifacts were corrected using independent component analysis (ICA, see Jung et al., 2000; mean number of components = 1). 7) The data without large artifacts were cleaned a last time using the threshold approach of step 4 (mean threshold absolute amplitude = 136.61 µV, SD = 45.34; mean threshold range = 212.35 µV, SD = 75.72; mean threshold variance = 2012.12 µV, SD = 2567.40), rejecting on average 3.91% of epochs (range 0.64% – 10.07). 8) Finally, bad channels were interpolated using the average of all neighbors (with regularization parameter λ = 1e-5).

Note that the total number of epochs rejected was small (average of 10.00% of epochs rejected, range = 3.99% - 25.75%) and that the proportion of remaining trials did not differ significantly between conditions (for all comparisons χ^2^(1) < 2.29, *p* > 0.78, Bonferroni corrected for multiple comparisons). The average remaining number of trials was 355.14 for the 1st triplet position, 355.40 for the 2nd, 357.14 for the 3rd, and 339.57 for random.

### Statistical analysis

#### Classification of learners

Participants were divided into two groups, “learners” and “other” using a conservative criterion. Learners were defined as individuals who scored on the behavioral offline test 22 or more out of 32. According to the binomial distribution this is the minimal score needed to present significantly above-chance learning at the individual level (with alpha = 0.05; see e.g., Bogaerts, Siegelman, & Frost, 2016; Siegelman, Bogaerts, & Frost, 2016). All participants that did not meet this criterion are considered “other”.

#### Spectral decomposition

All spectral estimates were based on time-frequency representations of power derived via a sliding-window fast Fourier transform (FFT). Spectral estimates were computed for frequencies between 1 Hz and 30 Hz on 0.5 s Hann tapered windows covering the pre-stimulus time interval between −0.8 s to 0 s (7 windows with a step size of 0.05 s). This pre-stimulus time window is largely uncontaminated by the visual evoked response for the preceding shape, which onsets 0.6 s prior to this window of interest. Each window was zero-padded to 1 s, providing a spectrally interpolated frequency resolution of 1 Hz. Hence we obtained a single pre-stimulus power spectral density estimate for each participant, channel, frequency and epoch by averaging the estimates for all 7 windows. Averaging in this way over each FFT window results in a more stable spectral estimate via Welch’s method with an overlap of 90 % (Welch, 1967).

#### Grand-average power spectrum

To calculate the grand-average power spectrum (shown in Figure 2) we averaged the spectral estimates for all trials, including positions 1, 2, 3 trials as well as random trials. We then averaged over subjects and electrodes. Spectral peaks were identified as maxima in the 1 – 30 Hz grand-averaged power spectra. Band-limited estimates were derived based on these peaks as the average of the maximum -+1 Hz, which we refer to as frequencies of interest (FOI). A simple normalization by frequency was applied dividing each power bin by 1/frequency.

**Figure 2.**
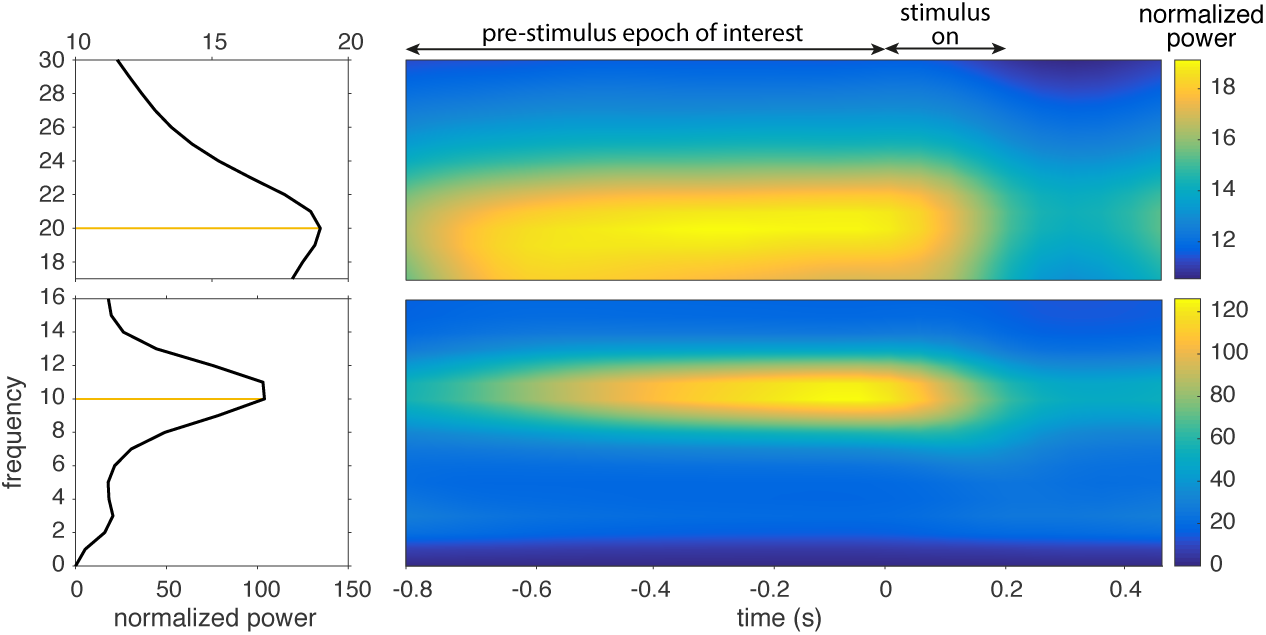
Left: peaks in the grand-average raw power spectrum of all pre-stimulus epochs within the structured and random stream. Yellow lines mark the peak frequency. Right: time-frequency plots for the grand-average. The black bar denotes the pre-stimulus epoch of interest.

#### Evolution of the spectral content of different pre-stimulus intervals

To investigate the evolution of spectral power over the course of the structured familiarization phase we computed average power per stimulus position for each channel and FOI for each of the 9 exposure periods (each containing 6 repetition blocks). To mitigate individual differences in overall raw EEG power, a normalization was performed such that the power for each position was expressed as the percentage of the average power over all trial types and channels, such that each trial type (positions 1, 2, 3, and random) contributed equally to this average.

To contrast the spectral power of the pre-stimulus windows prior to the three triplet position stimuli, taking into account the expected progression of learning over the course of the experiment, we used a cluster-based approach based on Monte-Carlo estimates of the null significance probabilities (Maris and Oostenveld, 2007). The average power over the 9 exposure periods of the structured stream were treated as the time-dimension in this analysis. For each FOI, a nonparametric permutation test clustered data samples of adjacent electrodes and time points (i.e., periods within the structured familiarization) simultaneously and compared the sum, of the descriptive statistic used, across each cluster. This approach is similar to the standard cluster-based analysis over electrode-time pairs within a trial, except here time is considered not at the trial level but at the larger scale of the 9 exposure periods of the structured familiarization stream (Maris and Oostenveld, 2007). The cluster-based tests were dependent samples and parameterized with a minimum number of neighboring channels of 3 and the cluster threshold t-value, or F-value corresponding to a p-value of 0.25. Note that the latter value does not affect the false alarm rate of the cluster test, it merely sets the threshold for considering a sample as a candidate member of a cluster (Maris and Oostenveld, 2007). This relatively low t-value functions to favor clusters with large spatio-temporal extent. All tests were based on distributions formed from 100000 permutations.

Note that we hypothesized spectral power differences as a result of assimilating the patterned structure of the stream; hence, for our initial analysis we only included the subset of individuals who demonstrated learning in the test phase (n=25). We performed a cluster-based permutation test over the three stimulus positions, computing an F-statistic, for all FOIs. The resulting p-values were Bonferroni corrected for the number of FOIs tested. Following the cluster-based F-test, post-hoc non-parametric paired t-tests were performed between each stimulus position, for each FOI that showed a significant difference for the cluster-based test. For these tests, the average power over subjects was computed for the set of all electrodes that were members of the significant cluster over the temporal extent of the cluster. Note, no more than one significant cluster was identified for any of the cluster-based tests. Each of the three t-tests was controlled for multiple comparisons using a max-based approach, where an omnibus null-distribution was constructed for the three t-tests based on the max-value over t-tests for each permutation (Nichols and Holmes, 2002; Maris and Oostenveld, 2007). For the beta band, we determined via the post-hoc t-tests that there was no difference between positions 2 and 3, while both these positions significantly differed from position 1 (see Figure 3). This motivated us to pool positions 2 and 3, and to perform a cluster-based t-test between this average and position 1. Expecting a greater level of statistical power and a desire to generalize results to all subjects despite their level of learning, this test was performed on all subjects, rather than only the ‘learner’ group.

**Figure 3.**
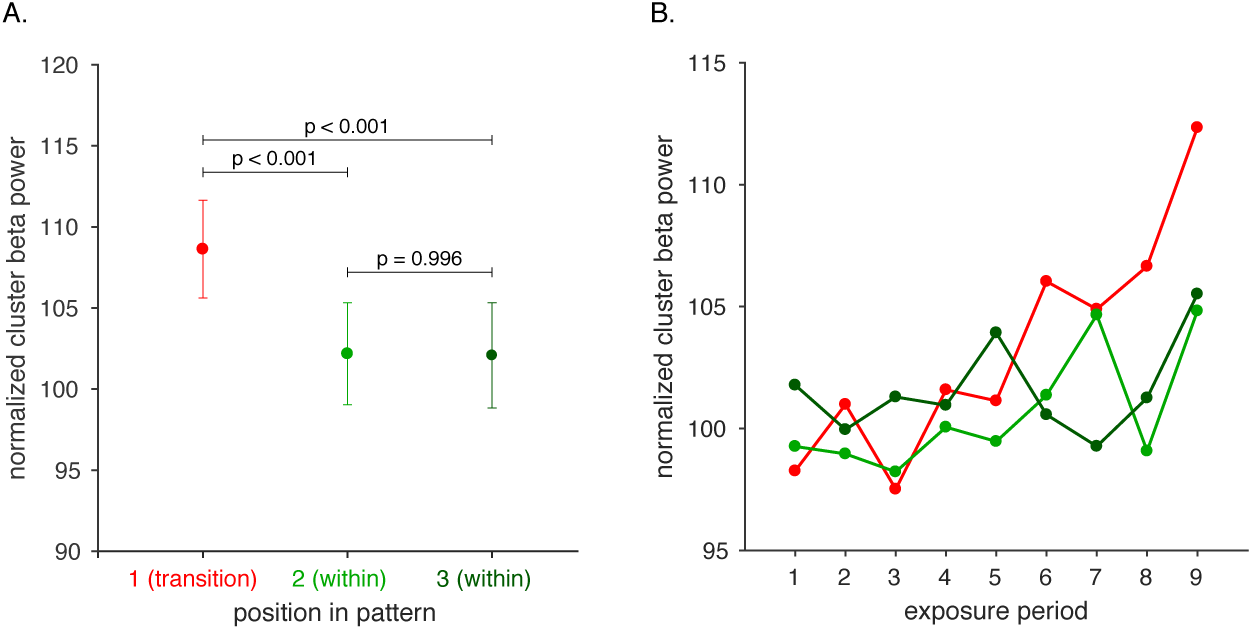
A. Average normalized beta power for the significant cluster. Error bars denote the standard error, p-values are corrected for multiple comparisons. B. Temporal evolution of the cluster beta power for each of the pre-stimulus intervals, across the structured familiarization stream. Exposure period 1 indicates the start of the structured familiarization stream, period 9 the end.

#### Brain-behavior correlation

The neural index of learning in the frequency domain (position 1 minus average of positions 2 and 3) was calculated per individual. This index was based on the average value over each electrode that was present in the significant cluster, across the temporal extent of the cluster. To assess the relation between our neural index of learning and offline test performance we use a Spearman rank correlation coefficient, which does not assume normality of the data, which helps to mitigate ceiling effect present in the post-test data.

#### Event related potentials

To evaluate if and how our frequency results relate to post-stimulus event related potential (ERP) differences between the triplet positions we also carried out analysis in the time-domain (Maris and Oostenveld, 2007). For this analysis the preprocessed data were band-pass filtered from 0.15 to 30 Hz, segmented into epochs ranging from stimulus onset to 0.5 s, and baseline corrected to a 200-ms pre-stimulus interval. Mean ERPs in the three triplet positions were calculated for each participant. To contrast the ERPs for the three triplet positions we took a cluster-based approach, clustering over channels and time points within the epoch using a nonparametric dependent samples T-statistic, with a minimum number of neighboring channels was 3. Note that an important advantage of this approach is that we were not restricting our analysis to a particular time window within the epoch or a particular electrode location.

### Data Sharing and Code Accessibility

The behavioral data reported in this paper, experimental materials and analysis code will be made available via the Open Science Framework (https://osf.io/). EEG data will be provided to any scientist upon request. This study has not been preregistered.

## Results

### Behavioral results

Average group performance on the offline test was 26.46/32 (SD = 6.41), which is significantly above chance (one-sample *t*-test comparing mean performance to a score of 16, corresponding to 50% chance level, *t*(34)= 9.6455, *p* < 0.0001). Based on a conservative individual criterion, 25 out of 35 participants were classified as learners.

### Spectral results

We identified two maxima in the average power spectrum across all participants: at 10 and 20 Hz (see 2). Subsequent analyses focused on these peak frequencies (FOIs) ± 1 Hz. We refer to the 19-21 Hz band as the beta range, and 9-11 Hz as alpha range.

#### Pre-stimulus beta power is dependent on stimulus position

Focusing on the data of learners, two cluster-based F-tests were performed in order to compare the spectral content of the pre-stimulus interval preceding the 1^st^, 2^nd^ and 3^rd^ stimuli in a triplet. We detected one significant positive cluster for the beta range indicating a difference between the three positions (Bonferroni corrected *p* = 0.01, see Figure 2-1), but none in the alpha range (Bonferroni corrected *p* = 0.11).

Post-hoc non-parametric paired t-tests for the significant cluster in the beta range revealed significantly higher power for the per-stimulus interval preceding a triplet transition (i.e., the interval before shapes in position 1), which by design was unpredictable (see Figure 3). No difference was found between the spectral power of the pre-stimulus interval preceding the two predicable positions 2 and 3, which were virtually identical. Figure 3 also shows the evolution of average cluster beta power over the 9 exposure periods. What we observe is a gradual divergence of the beta power at pattern transitions relative to the beta power within patterns. An ANOVA with position and exposure period as repeated measures factors indicates a significant effect of position (*F*(2,48) = 4.20, *p* = 0.021) as well as a significant interaction between position and exposure period (*F*(16,384) = 2.10, *p* = 0.008).

#### Beta power within versus between pattern transitions

Given the difference between the pre-stimulus beta-frequency power of the unpredictable 1^st^ shape and the predictable shapes of positions 2 and 3, we subsequently focused on the contrast of position 1 versus the average of positions 2 and 3. This contrast revealed –in line with the results reported above– a significant cluster (*p* = 0.0038). As this contrast served as an individual index of learning, which we correlated with offline test scores, we performed the cluster-based t-test also including all participants. Figure 4 illustrates the significant cluster (*p* = 0.003) we observed for the entire sample, revealing a broad central scalp distribution across exposure periods 4 to 9.

**Figure 4.**
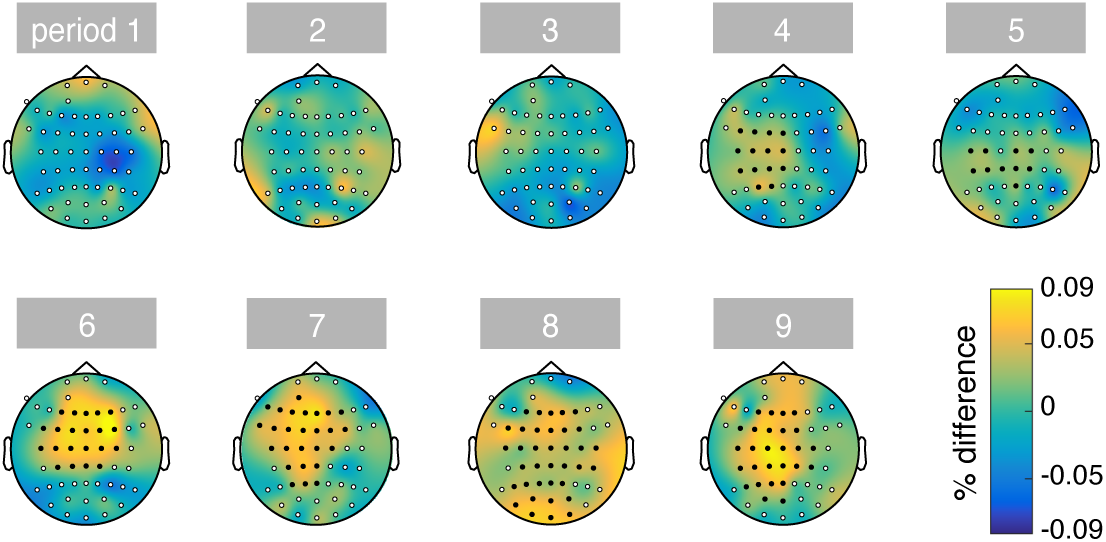
Temporal evolution of the topography of the difference between pre-stimulus beta-band power for within versus between triplet transitions. Period 1 indicates the start of the structured familiarization stream, period 9 the end. Electrodes that are part of the significant cluster are filled black.

**Figure 5.**
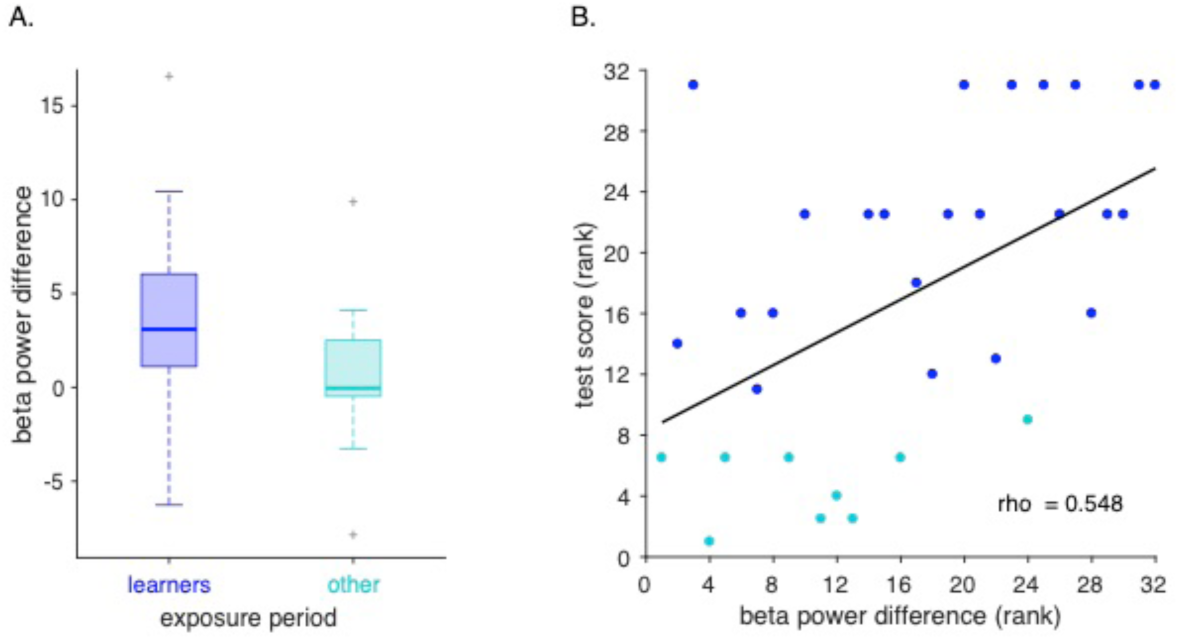
A. Boxplot summarizing the beta power difference for participants classified as learners versus others. On each box, the central line corresponds the median and bottom and top edges of the box indicate percentiles 25th and 75th. Dashed whiskers extend to the most extreme data points not considering outliers. Outliers are represented by a ‘+’ symbol. B. Relation between the size of the beta power modulation and behavioral test scores.

#### Correlation between the Beta indices of learning and offline test scores

We next validated our neural signature of learning by investigating whether the difference in beta power for within versus between pattern transitions covaries with the behavioral offline measure of learning across individuals. We observe a strong positive relation between these two indices of learning (*Spearman’s rho* = 0.55, *p* = 0.0007).

### ERP results

No significant clusters were observed when comparing the grand average ERPs for either position 1 versus the average of 2 and 3, position 1 versus 2, or position 1 versus 3. When calculating the ERP based on exposure periods 4 to 9 of the experiment only, we again find no significant clusters for either contrast. This suggests that our frequency finding in the pre-stimulus interval occurs in the absence of any significant modulation of ERPs due to the learning of statistical structure (smallest corrected p-value = 0.19).

## Discussion

In the current study we identified a neural signature of visual SL, operationalized as the extraction of triplet patterns embedded in a continuous sequence of abstract shapes given differences in transitional probabilities. This signature comprised increased beta band activity in the interval leading up to unpredictable shapes in the stream, thus, this effect was concurrent with triplet boundaries. Importantly, our results indicate that the differential beta band power prior to unpredictable shapes increased steadily over the course of repeated exposure to the patterns. Moreover, looking at individual differences in the magnitude of this effect, we show that the differential pre-stimulus beta power is highly predictive of performance in an offline behavioral test of pattern recognition, which is the learning measure used by the vast majority of SL studies (see Siegelman, Bogaerts, Christiansen, & Frost, 2017). This leads us to conclude that our proposed spectral signature is tracking the learning process of segmenting the continuous stream, and thus provides a valid online assessment of SL performance.

The identification of a neural signature of visual SL offers novel insights regarding the mechanisms underlying SL. The increased beta power at triplet transitions point to two theoretical possibilities. A first possibility is that such beta-band activity reflects anticipation of uncertainty (i.e., higher entropy) once the statistical structure of the recurrent patterns in the stream is assimilated. Moreover, in addition to uncertainty, the upcoming item, consisting of the first position of a new triplet is highly informative of the identity of this triplet. Thus, the maximal preparation of processing resources may be expected prior to the first item of a triplet. This interpretation concurs with the proposed functional role of beta oscillations for attentional top-down regulation in the literature (see Engel & Fries, 2010 and Bastos & Friston, 2012, for a review). On a speculative note, the direction of our effect (i.e., *more* beta power prior to pattern transitions relative to transitions within a visual pattern) might reflect a distributed, global beta network state in anticipation of high uncertainty and a focused, local state in anticipation of predictable events. Linking back to other neurophysiological findings in the literature, the assumption is that intracranial recordings pick up on the latter state.

A second possibility is that the increased beta oscillations reflect post-processing of the now completed triplet. If one regards a learned triplet as a cognitive set that is the target of learning, it is possible that the interval between triplet transitions is used to actively maintain this set in memory. Indeed such functional role has been hypothesized for beta-band activity (Engel and Fries, 2010). This is further evident in findings reporting elevated beta-band activity in the delay phase of working memory tasks (Deiber et al., 2007; Siegel et al., 2009; Salazar et al., 2012). Similarly, if learning results in the grouping of stimuli into chunks and these chunks are what is stored in memory (Orbán et al., 2008; Perruchet, 2018), the oscillatory beta signal could reflect the process of memory encoding or strengthening (e.g., Berke, Hetrick, Breck, & Greene, 2008). However, note that the learning that occurs in the current SL task is incidental (different from typical working memory tasks).

Our finding that we can track learning during passive exposure to regularities without monitoring overt responses, has important implications for assessing learning. The caveats involved in assessing learning through post-familiarization two-alternative-forced-choice questions have been discussed in length (see for example, Siegelman et al., 2017). Our findings offer then an online neurobiological signature of learning that reveals a learning trajectory over time, without a need for behavioral testing.

In conclusion, our findings reveal a neural signature of visual regularity learning: elevated beta band activity at pattern transitions. This signature tracks the segmentation process during pattern exposure and is highly predictive for the behavioral learning outcome. Whether the heightened beta-band activity reflects the anticipation of a novel upcoming pattern or rather post-processing of the completed pattern requires additional investigation, aiming to unravel the possible functional role(s) of beta-band oscillations in regularity learning.

The authors report no financial interests or conflicts of interest.

## Acknowledgments

This paper was supported by the ERC Advanced grant awarded to Ram Frost (project 692502-L2STAT) and Marie Skłodowska-Curie Grant No. 743528 (IF-EF, European Union’s Horizon 2020 Research and Innovation Programme) awarded to Louisa Bogaerts. We wish to thank Limor Shedlesky, Cassi Gewer and Michelle Schechter for their help with subject recruitment and data collection. L.B.’s current affiliation is Free University Amsterdam.

## Extended data

**Figure 2-1.**
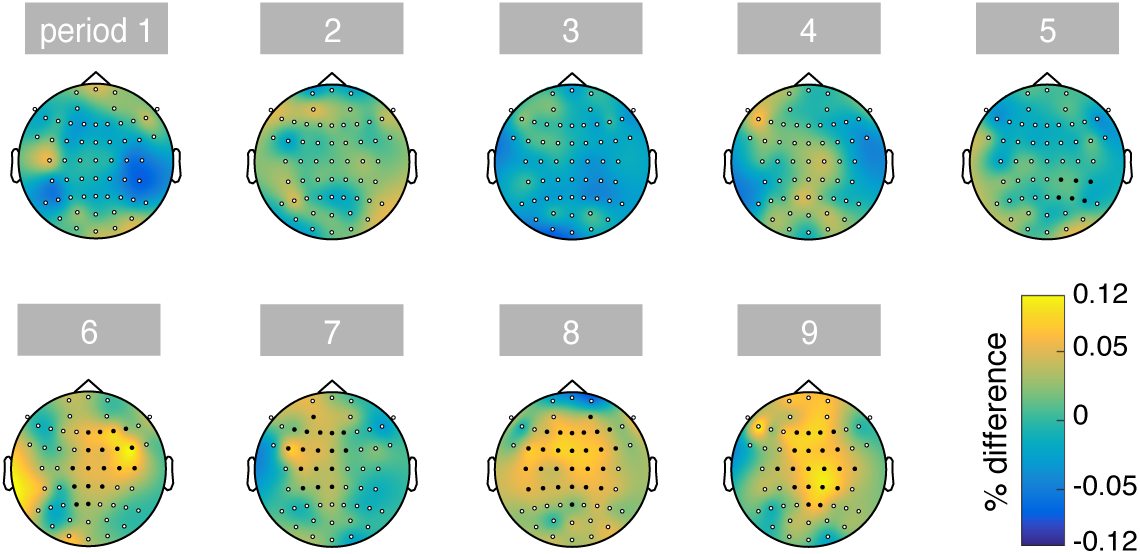
Temporal evolution of the topography of the F-cluster comparing pre-stimulus beta-band power prior to shapes in position 1, 2 and 3. Period 1 indicates the start of the structured familiarization stream, period 9 the end. Electrodes that are part of the significant cluster are filled black.

## Notes

### Competing Interest Statement

The authors have declared no competing interest.

